# Unraveling Coinfection Dynamics into 100 Whole Genome of Diarrheal Pathogens: A Genome-to-Systems Biology Approach with *Plesiomonas shigelloides*

**DOI:** 10.1101/2023.11.24.568624

**Authors:** Mohammad Uzzal Hossain, A.B.Z Naimur Rahman, Md. Shahadat Hossain, Shajib Dey, Zeshan Mahmud Chowdhury, Arittra Bhattacharjee, Ishtiaque Ahammad, Md. Imran Ahmed, Khandokar Fahmida Sultana, Abu Hashem, Keshob Chandra Das, Chaman Ara Keya, Md. Salimullah

## Abstract

Diarrhea is the second leading cause of mortality among infants under the age of five. One of the main causes of this disease is multipathogenic infections, which can make the conditions of patients even worse. *Plesiomonas shigelloides* (*P. shigelloides*) is one of the pathogenic bacteria that contributes to the pathophysiology of diarrhea and may be implicated in coinfection with other diarrheal pathogens. Therefore, the purpose of this study is to investigate the hypothetical proteins to explore the genetic insights of *P. shigelloides* and its relationships with common diarrheal diseases. For this reason, we used 16S rRNA sequencing together with several biochemical tests to identify the bacteria that we isolated from diarrheal patients (8 years). Afterwards, the whole genome of *P. shigelloides* was sequenced, assembled and annotated in order to obtain the genomic insights of *P. shigelloides*. In addition, the common virulence genes of ten (10) common diarrhea-causing bacteria were identified from 100 whole genome sequences. Finally, the system biology approach was applied to predict the coinfection pattern between *P. shigelloides* and the virulence genes of 10 bacteria. The complete genome sequencing analysis of this bacterium revealed 899 hypothetical proteins from which 33 hypothetical proteins shared the clusters with the 109 virulence genes of 10 distinct diarrheal pathogens, forming a strong interaction based on biological processes, molecular functions, subcellular localization, or pathways. All diarrhea causing bacteria were found to have *P. shigelloides* microbial interactions; however, *V. cholerae* exhibited the strongest relationships, while *C. difficile* showed the weakest. The results of this investigation clearly imply that *P. shigelloides* shares a coinfection pattern with other bacteria that cause diarrhea. Finally, the findings from the complete genome provide new avenues for researchers to pursue their investigation of the pathophysiology of diarrhea.

## 1. Introduction

Diarrhea is a disease of the gastrointestinal tract characterized by frequent, loose, and watery bowel movements. Globally, 1.5 million deaths and nearly 1.7 billion diarrheal cases occurred every year [1]. It is also the second leading cause of death in children <5 years old and is responsible for the death of more than 525,000 children every year worldwide [2]. The Global Enteric Multicenter Study (GEMS) study says that diarrheal disease is a major threat to human health and is still the top cause of mortality and morbidity around the world, especially in developing countries like sub-Saharan Africa and south Asia [3, 4]. In Bangladesh, diarrheal disease remains one of the leading causes of death and the most prevalent reason for hospitalization [5-7]. The overall prevalence of diarrhea was found to be 5.71% among children <5 years old in Bangladesh [8].

Diarrhea is caused by pathogenic microorganisms including viruses, bacteria and parasites [1]. Although, viruses cause the majority of diarrheal disease, approximately 15-20% of diarrhea is caused by bacteria [9]. In developing countries, bacterial pathogens and parasites are the leading cause of infectious diarrhea [10]. The common bacterial pathogens that cause diarrheal disease are *Salmonella spp, Shigella spp, Vibrio cholerae, Vibrio parahaemolyticus*, Enteroinvasive *E. coli* (EIEC), Enterotoxigenic *E. coli* (ETEC), *Campylobacter jejuni*, and *Clostridium difficile* [11, 12]. Apart from these, there is another bacterium *Plesiomonas shigelloides*, that is currently very concerned to public health [13]. This bacterium has been ignored for many years, but in recent years it has begun to be recognized as a true enteropathogen and disease-causing microorganisms in both gastrointestinal and extraintestinal complications [14]. *Plesiomonas shigelloides* is also responsible for 11 outbreaks among five countries [13] and has some strong evidence of coinfection among other diarrheal pathogens [15]. Coinfections may result in more severe or prolonged symptoms of diarrhea, fever, abdominal pain and vomiting. It may interact with other diarrheal pathogens by competing for nutrients, altering the intestinal environment, modulating the host immune response or enhancing the virulence of other pathogens [15, 16]. So, research on genomic insights of *Plesiomonas shigelloides* could bring new hope to treatment of diarrheal disease.

In previously reported on diarrheal treatment and medication, all of the research were based on known essential proteins and virulence genes of diarrheal pathogen. Glucose-electrolyte oral rehydration therapy (ORT) is the most common treatment for diarrheal disease that can reduce acute mortality from dehydration [17] and only in cases of serious bloody diarrhea or dysentery antibiotic medication are advised [18]. However, some diarrheal pathogens have shown antimicrobial resistance through mutation and alteration of proteins that are antimicrobial targets and that’s why this treatment isn’t more effective against diarrheal disease [19]. This circumstance necessitates to focused on other virulence genes along with essential proteins for find out more effective therapeutic targets for the treatment of diarrheal disease.

Study on the hypothetical protein (HP) of *Plesiomonas sp*. could be an effective therapeutic target for this purpose. In the realm of molecular biology and bioinformatics, a hypothetical protein is a protein that has been predicted to exist based on the presence of a corresponding gene in a genome or transcriptome, but for which there is limited or no experimental evidence to confirm its existence or function. In the case of bacteria, 25-50% of the genome comprises HPs, which may have roles in disease and therapeutic targets because they are involved in the pathogenesis of a disease or the regulation of a biological pathway [20]. Hypothetical proteins also have roles in microbial interactions by affecting the expression, regulation or function of other proteins in the host or the microbiome [21, 22]. Some methods to predict the function of hypothetical proteins using big data and machine learning are based on clustering the proteins based on their sequence similarity or interaction patterns [20, 22].

To investigate of genomic data of a bacterium, it is necessary to have enough data and studies on this bacterium. Unfortunately, there are few studies are available on *Plesiomonas shigelloides* in Bangladesh, and the NCBI database doesn’t have any WGS sequence data from Bangladeshi cases. Though *Plesiomonas* have an important role in diarrheal disease and have strong evidence of coinfection with other diarrheal pathogens, genomic studies on this bacterium are important for the treatment of diarrheal disease.

For this purpose, our study has been conducted with genomic insights of *Plesiomonas shigelloides* and to find out the microbial interaction of hypothetical proteins of this bacterium with other diarrheal pathogens.

## 2. Methods and Materials

### A. Clinical isolates of diarrheal pathogen

#### 2.1. Clinical sample collection

A total of 11 stool samples were collected from diarrheagenic infants (≤8 years) from Dhaka Shishu Hospital, Dhaka. The sample metadata presented in **Supplementary Table 1** comprises demographic information regarding the age, origin, and gender of the patients. All the samples were collected aseptically using sterile specimen containers and transported to the Microbial Biotechnology laboratory within 2 hours for further isolation and identification studies.

#### 2.2. Isolation and Identification of Bacteria

A loopful of collected stool samples was inoculated in 9 ml of Tryptic soy broth and incubated at 37°C for 18 h for enrichment. After enrichment, a loopful of inoculums was then streaked on MacConkey agar (Oxoid, UK) and Salmonella-Shigella agar (Oxoid, UK) plates and incubated at 37°C for 24 h. Non-lactose fermenters and transparent or colorless colonies from MacConkey plates and colorless colonies from Salmonella-Shigella agar plates were further streaked on Salmonella-Shigella agar plates and incubated at 37°C for 24 h. Suspected colonies of *Plesiomonas* from selective medium were picked and streaked onto Nutrient agar plates and incubated at 37º C for 18-24 h for its biochemical characterization. Typical colonies were selected from Nutrient agar plates and were subjected to slide preparation, Gram staining, microscopic observation, and biochemical tests. Pure colonies were identified by biochemical tests, triple sugar iron agar (TSI), motility test, urease test, indole test, citrate test, and oxidase test.

#### 2.3. Bacterial genomic DNA extraction and purification

Bacterial genomic DNA extraction and purification were done by using Thermo Scientific GeneJET Genomic DNA Purification Kit and their extraction protocol. The GeneJET™ Genomic DNA Purification Kit is designed for rapid and efficient purification of high-quality genomic DNA from bacteria. The kit utilizes silica-based membrane technology in the form of a convenient spin column. After purification, DNA concentration and purity were measured by using a NanoDrop™ 2000/2000c Spectrophotometer.

#### 2.4. Amplicon generation, sanger sequencing and phylogeny analysis

To identify bacteria, their 16S rRNA genes were amplified by PCR using primers 27F (5′-AGAGTTTGATCCTGGCTCAG-3′) and 1492R (5′-TACGGYTACCTTGTTACGACTT-3′) [23], which amplified nearly the entire 16S rRNA gene [23]. The amplified PCR products were resolved by agarose gel electrophoresis. The DNA sequence of both strands was determined by using sanger sequencing protocol from Molecular Biotechnology Division, National Institute of Biotechnology, Bangladesh with an Applied Biosystems 3130 Genetic Analyzer and was analyzed with Sequencing Analysis 5.3.1 software (Applied Biosystems).

The sequencing results were analyzed by Bioedit sequence analysis software and performed BLAST (optimized for highly similar sequences) against the NCBI database to determine the species. The neighbor-joining method was used to construct phylogenetic trees using Molecular Evolutionary Genetics Analysis (MEGA) 7.0 software. A bootstrap test was performed with 1000 repetitions.

### B. Genomic insights of diarrheal pathogen

#### 2.5. Next Generation Sequencing (NGS) and data preprocessing of the clinical isolate

The Sanger sequence identified and confirmed bacterial isolates were sequenced using the MiSeq system of Illumina sequencing platform at the center for next-generation sequencing facility at the National Institute of Biotechnology (NIB). First, the extracted genomic DNA was digested into 400–550 bp fragments by using the M220 Focused-ultrasonicator (Covaris Ltd, Brighton, UK). The sequencing library was constructed using the Nextera XT DNA Library LT kit (Illumina, San Diego, CA, USA), according to the manufacturer’s protocols. WGS was performed on an Illumina MiSeq system (Illumina) with a 300 bp paired-end reads sequencing kit (MiSeq Reagent Kit v3; Illumina). The raw data from the MiSeq instrument in the gz compressed FASTQ format were analyzed with several bioinformatics tools in the Linux operating system. FastQC and MultiQC tools were used to perform quality control checks on raw sequence data and the trimmomatic tool was used for filtering low-quality reads and removing adapter sequences from reads [24].

#### 2.6. Assembly, Annotation, and Identification of the hypothetical proteins

The genome assembly was then carried out using the Unicycler [25], which uses SPAdes (v3.6.2 or later) to construct a De Bruijn graph assembly using a wide range of k-mer sizes. When assembly was completed, the quality of the assembly was checked with the tool QUAST. QUAST provides detailed information about contigs, such as N50 length which is defined as the shortest sequence length at 50 % of the genome. The contigs of the assembly were then annotated for protein-coding sequences, tRNA, and small RNAs using the tool Prokka [26]. The annotated genome is then explored briefly using Artemis [27] to gain proficiency with the tool. The hypothetical protein sequences were then separated from the annotated file for further study.

#### 2.7. Retrieval of 100 bacterial whole genome sequencing data

100 bacterial WGS sequencing data (10 *Shigella dysenteriae*, 10 *Shigella flexneri*, 10 *Vibrio cholerae*, 10 *Vibrio parahaemolyticus*, 10 *Clostridium difficile*, 10 *Salmonella enterica* serovar Typhimurium, 10 *Salmonella enterica* serovar Enteritidis, 10 *Campylobacter jejuni*, 10 Enteroinvasive *Escherichia coli* and 10 Enterotoxigenic *Escherichia coli)* from the Illumina platform was retrieved from the NCBI Sequence Read Archive (SRA) database, and these all bacteria were collected and sequenced from stool samples of diarrheal patients in the Asian region (**Supplementary Table 2**). The sequence reads were downloaded by using the SRA-Toolkit tool as FASTQ files.

#### 2.8. Assembly of 100 WGS sequencing data and screening of virulence genes

All the sequencing data were assembled with the hybrid de novo assembler Unicycler. Genes related to virulence were detected using ABRicate [28] by means of the Virulence Factor Database (VFDB) [29]. Only hits with >90% identity and >60% target coverage are retained. The common virulence genes were identified for each bacterium.

### C. System biology approach to detect the microbial interactions

#### 2.9. Protein-Protein interaction between hypothetical protein of *P. shigelloides* and virulence genes

In this step, we used the STRING database to identify microbial interactions between *Plesiomonas shigelloides* and 10 different diarrheal pathogens. STRING is a database of known and predicted protein-protein interactions (PPI) [30]. The hypothetical protein sequences were employed to established PPI network with the virulence genes of 10 distinct bacteria. Using the degree distribution of nodes, the network of diarrheal pathogen and *P. shigelloides* was analyzed to identify highly connected proteins. Then we clustered the network into one, two or three groups using the K-means clustering method through the STRING web server [30]. The network files were then visualized using the Cytoscape software, and interaction analysis was used to guess which nodes were connected. The prediction model of STRING was used to find the gene ontology of biological function, metabolic process, and KEGG pathways. Different layout choices were used to see the cluster networks, and high-quality network pictures were made to see them in more depth.

## 3. Results

### 3.1. Morphological Identification

Among 11 samples, one sample (H-11) has been shown *Plesiomonas shigelloides* like colonies. The bacterium grew well on MacConkey agar and Salmonella-Shigella agar after 18 hours of incubation at 37° C. Colonies of the isolates were non-lactose fermenting, round, beige opaque, moist, and slightly raised on MacConkey agar plates **(Figure 1A)**. In the Salmonella-Shigella agar medium, colonies were transparent or colorless, circular, entire, and slightly raised (**Figure 1B**). Gram staining results were microscopically observed. The bacterial cells stained red and appeared as short rods, which were arranged singly or in pairs, indicating that the strain was gram negative **(Figure 1C)**. The biochemical characteristics of *Plesiomonas shigelloides* are provided in **Table 1**. *Plesiomonas* is oxidase, catalase and motility positive, and alkaline over acid with no gas or hydrogen sulphide in TSI (**Supplementary Figure 1**).

**Figure 1:**
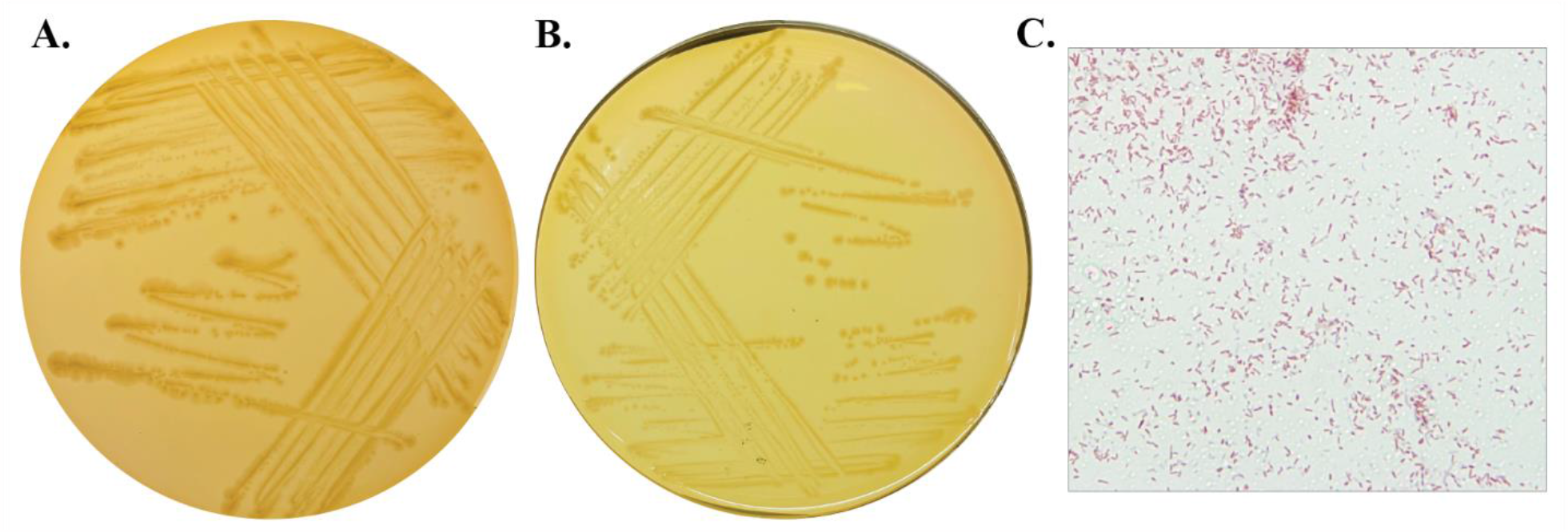
Colonies of *P. shigelloides* on (A) MacConkey agar and (B) Salmonella-Shigella agar plates, and (C) microscopic features of *P. shigelloides*.

**Table 1:** Biochemical results of *P. shigelloides*.

| Sample ID | MIU medium |  |  | TSI medium |  |  |  | Citrate | Oxidase | Catalase |
| --- | --- | --- | --- | --- | --- | --- | --- | --- | --- | --- |
|  | Motility | Indole | Urea | Slant | Butt | H<sub>2</sub>S | Gas |  |  |  |
| H-11 | + | + | - | Alkaline (Red) | Acidic (Yellow) | - | - | - | + | + |

### 3.2. PCR and Sequence Analysis

PCR products of the *Plesiomonas* were 1400-1500 bp in range while amplified using 27F and 1492R primers **(Figure 2A)**. The amplicons were then purified and sequenced using Sanger sequencing. NCBI BLAST was used for homology analysis based on the 16S rRNA gene sequences of the strain; the results revealed that the strain was 100% similar to NR_044827.1 *Plesiomonas shigelloides* strain. Phylogenetic analysis was performed after screening sequences with high similarity in terms of sequence alignment; the results (**Figure 2B**) revealed that 16S rRNA gene sequences of *P. shigelloides* formed a single cluster. Therefore, based on its morphological, physiological, biochemical, and molecular characteristics, the strain was identified to be *P. shigelloides*.

**Figure 2:**
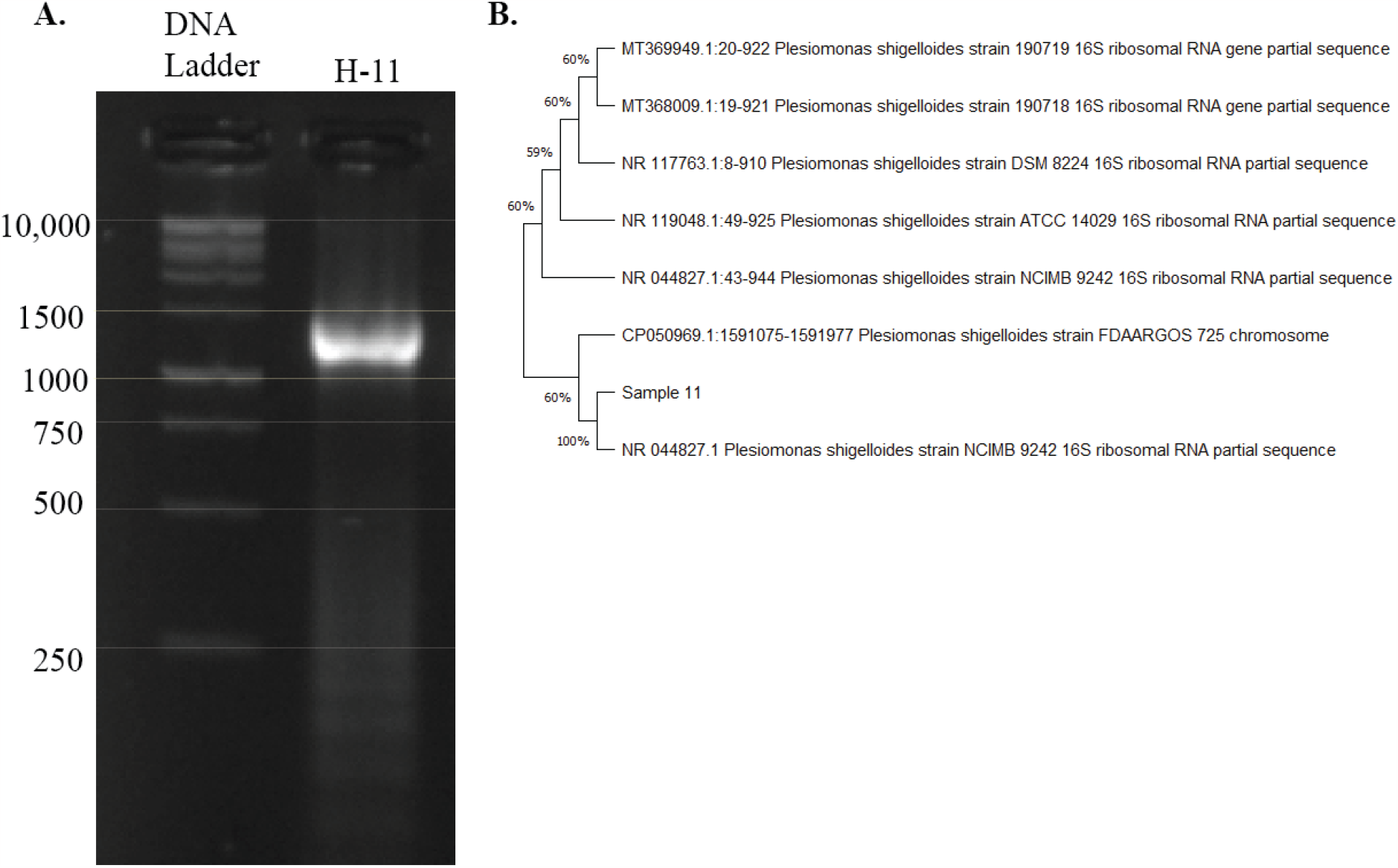
(A)Agarose gel electrophoresis (1.5%) of PCR product using 27F and 1492R primers. (B) Phylogenetic tree based on the partial 16S rRNA gene sequences; the algorithm used to construct the tree was the unweighted pair group method using averages (UPGMA).

### 3.3. Genome Assembly and Annotation of *Plesiomonas shigelloides*

The assembly results indicated no presence of contaminating sequences from other organisms and have shown 100% identity with *Plesiomonas shigelloides*. The Unicycler Assembler assembled 50 contigs with a total genome size 3617790 base pairs having 51.92% GC contents. The results of QUAST indicated there are no mismatches in this assembly and the largest contig was 461416 bp. The QUAST reports (**Supplementary Figure 1**) have shown that N50 was 299831 bp and N90 was 97454 bp.

Genome annotation was performed by Prokka tools and assembled genome of *Plesiomonas shigelloides* have 3099 coding DNA sequences (CDSs), 4 rRNAs, 99 tRNAs, 1 tmRNA, and 899 hypothetical proteins. The hypothetical proteins were mapped to their functional assignment using the STRING server. Among 899 hypothetical proteins, 33 hypothetical proteins were selected for the study based on STRING mapping (**Supplementary Table 3**). A complete graphical display of the distribution of the genome annotations is provided in **Figure 3**.

**Figure 3:**
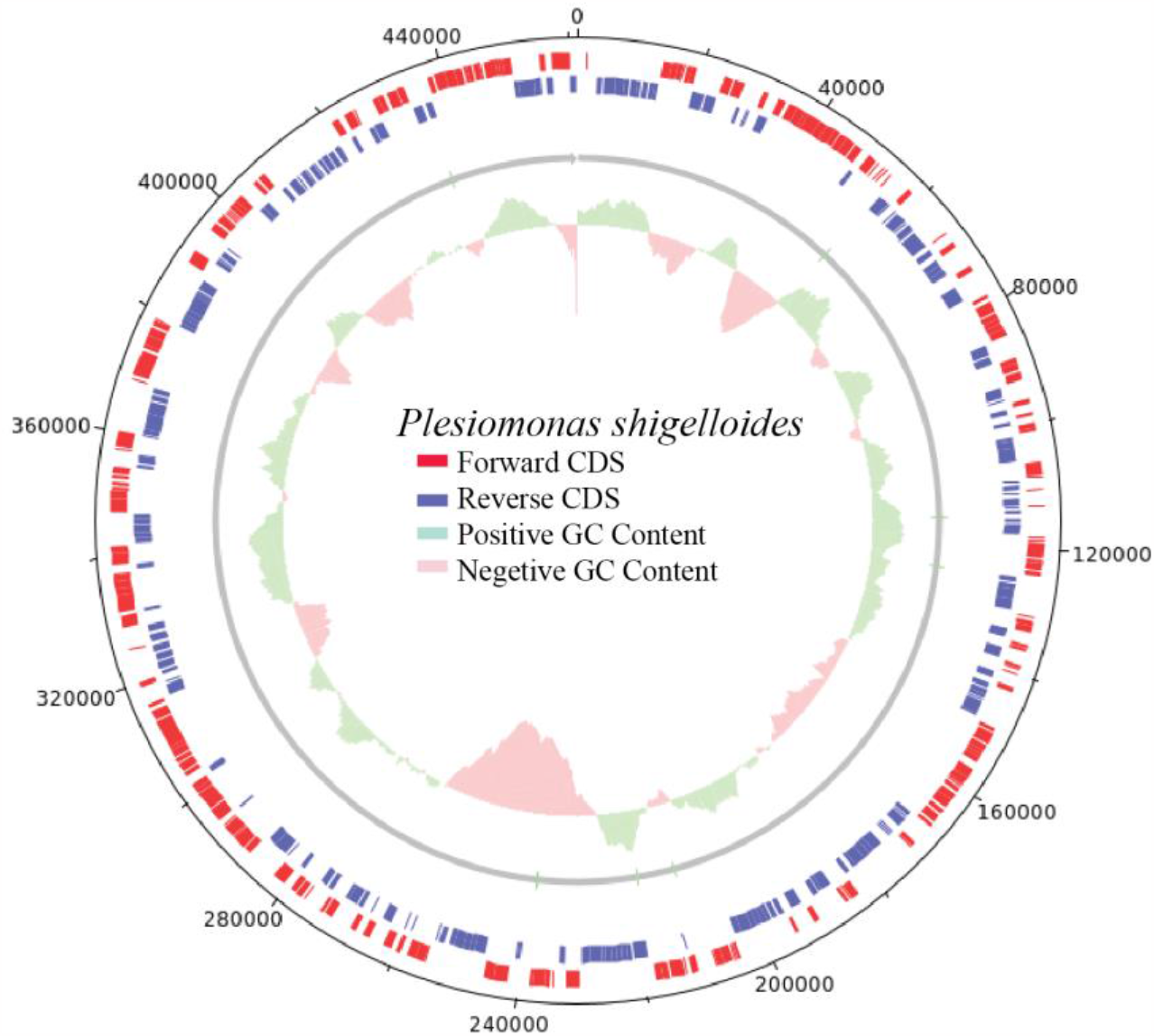
Complete genome plot of *Plesiomonas shigelloides*; Red line represents the forward coding DNA sequences (CDSs) and blue line represents reverse coding DNA sequences, green and pink colors represent the GC content value.

### 3.4. Nucleotide sequence accession numbers

The draft genome sequences for *Plesiomonas shigelloides* NIB001 have been deposited at NCBI GenBank under the accession no. JAVRQE000000000 (Version: JAVRQE000000000.1, BioProject: PRJNA1017991, SRA ID: SRR26106882).

### 3.5. Identification of Virulence genes from 100 WGS of diarrheal pathogens

After the assembly of 100 bacterial WGS sequencing data, we identify total 539 virulence genes by using the ABRicate tools that use Virulence Factor Database (VFDB) as a reference database (**Supplementary Table 4**).

### 3.6. Microbial interactions of *P. shigelloides* with 10 diarrheal pathogens

In this study, we used 10 *Shigella dysenteriae* and 10 *Shigella flexneri* WGS sequencing data to find out the virulence genes from vfdb database. Among these virulence genes, we filtered 41 common virulence genes between these two species (**Supplementary Table 4**). These proteins established a Protein-Protein Interaction (PPI) network with 3 hypothetical proteins (NIB_SUN.1, NIB_SUN.2 and NIB_SUN.3) of *P. shigelloides*. A cluster was established between three hypothetical proteins with fur, phoP and rpoS virulence proteins of *Shigella spp*. and the interactions were based on gene neighborhood and co-expression (**Figure 4A**).

**Figure 4:**
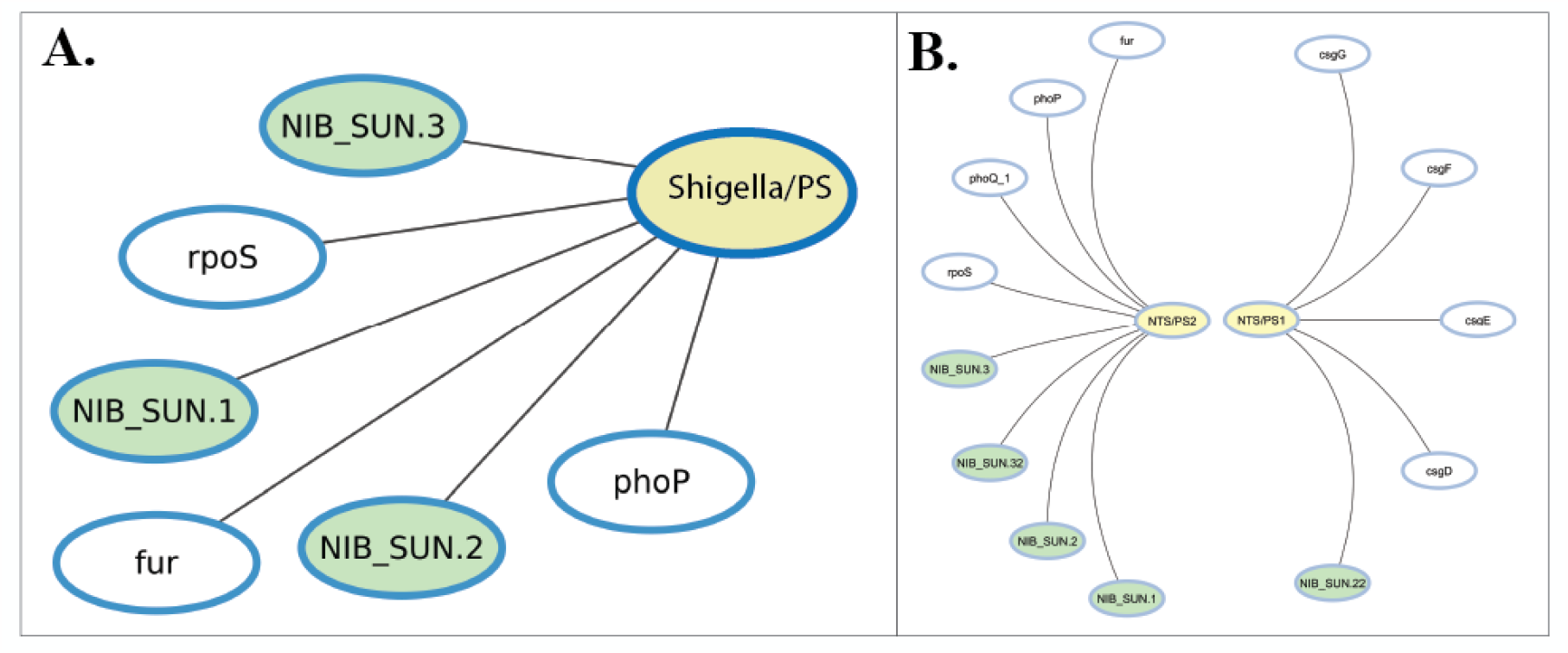
Microbial interaction of (A) *Shigella spp*. and (B) *Salmonella spp*. virulence genes with *Plesiomonas shigelloides*. Shigella/PS, NTS/PS1 and NTS/PS2 are cluster name; Green nodes: Hypothetical proteins of *P. shigelloides*, and White nodes: virulence genes of (A) *Shigella spp*. and (B) *Salmonella spp*.

131 common virulence genes were selected from the analysis of non-typhoidal *Salmonella* (*Salmonella enterica* serovar Typhimurium and *Salmonella enterica* serovar Enteritidis) WGS sequencing data (**Supplementary Table 4**). Two clusters were established between non-typhoidal *Salmonella spp*. (NTS) and highly interconnected hypothetical proteins of *P. shigelloides*, resulting in an overall network comprising 13 nodes and 15 edges (**Figure 4B)**. NTS/PS1 cluster have 5 nodes and 8 edges of virulence genes (csgD, csgE, csgF, csgG) and 1 hypothetical protein (NIB_SUN.22). This cluster associated with the biological process of pilus assembly and protein transport, molecular function of identical protein binding and subcellular localization of cell envelope. NTS/PS2 cluster contained network with 8 nodes and 7 edges of 4 virulence genes (fur, phoP, phoQ and rpoS) and 4 hypothetical proteins (NIB_SUN.1, NIB_SUN.2, NIB_SUN.3 and NIB_SUN.32) based on gene neighborhood, gene fusions, gene co-occurrence, co-expression and protein homology.

A total of 40 virulence genes were chosen based on the examination of whole-genome sequencing (WGS) data obtained from NCBI Sequence Read Archive (SRA) database (**Supplementary Table 3**). This selection included 10 Enteroinvasive *Escherichia coli* and 10 Enterotoxigenic *Escherichia coli*. A cluster was formed including three hypothetical proteins of *P. shigelloides* with three virulence genes of *E. coli* (**Figure 5**). The establishment of this cluster was determined by the analysis of gene neighborhood and co-expression.

**Figure 5:**
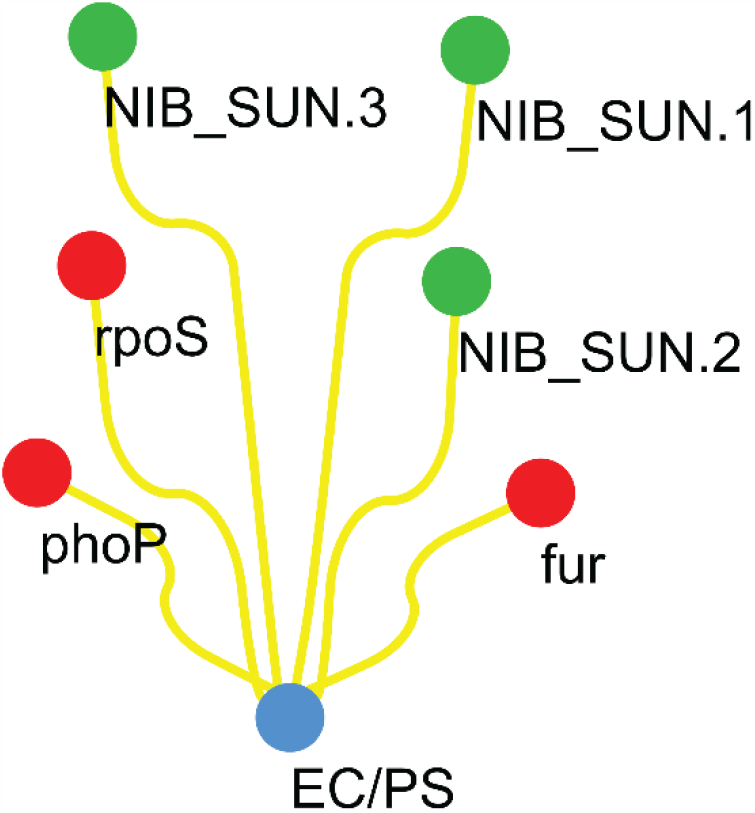
Microbial interaction of *Escherichia coli* virulence genes with *Plesiomonas shigelloides*. EC/PS cluster name; Green nodes: Hypothetical proteins of *P. shigelloides*, and Red nodes: virulence genes of *Escherichia coli*.

In the case of *C. jejuni*, three clusters (CJ/PS1, CJ/PS2, CJ/PS3) were established between *C. jejuni* and highly interconnected hypothetical proteins of *P. shigelloides*. The CJ/PS1 cluster consisted of 13 nodes and 8 edges, with 9 virulence genes of C. jejuni and 4 hypothetical proteins of *P. shigelloides* (**Figure 6)**. Carbohydrate metabolic process, lipopolysaccharide biosynthesis, are all related to this cluster. Meanwhile, the CJ/PS2 cluster encompassed 42 protein nodes and 549 edges of 10 hypothetical proteins and 32 virulence genes (**Figure 6)**. This cluster is related to several biological processes, especially chemotaxis, protein transport and response to external stimulus. It also related to molecular function, KEGG pathways, and subcellular localizations of flagellum. In CJ/PS1 cluster, there were 8 nodes and 15 edges, with 6 hypothetical proteins and 2 virulence genes (**Figure 6)**. This cluster is connected to a number of biological functions, particularly protein transmembrane transport and protein secretion.

**Figure 6:**
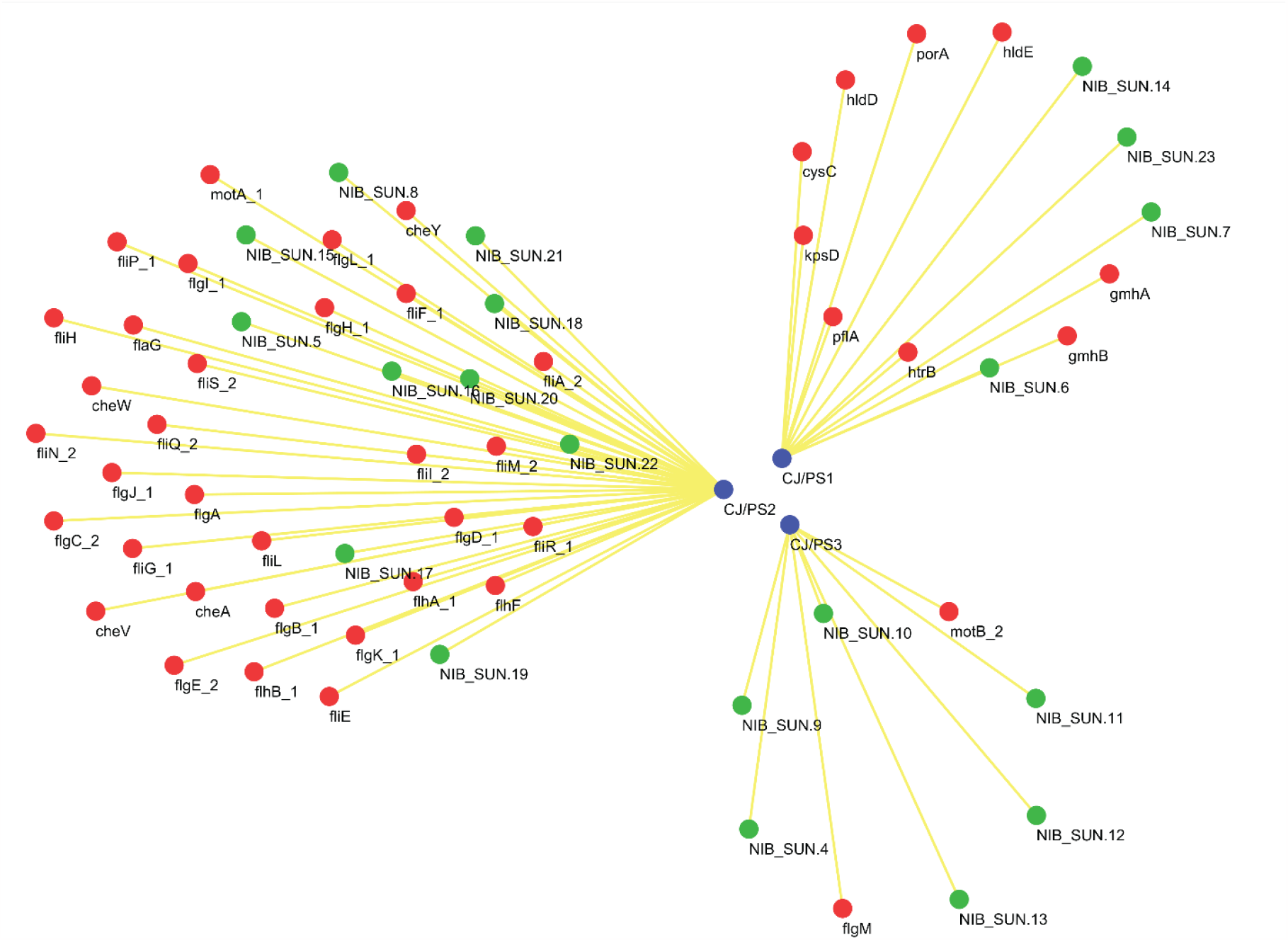
Microbial interaction of *Campylobacter jejuni* virulence genes with *Plesiomonas shigelloides*. CJ/PS1, CJ/PS2 and CJ/PS3 cluster name; Green nodes: Hypothetical proteins of *P. shigelloides*, and Red nodes: virulence genes of *Campylobacter jejuni*.

Virulence genes of *V. cholerae* established three distinct network clusters (VC/PS1, VC/PS2 and VC/PS3) when interacting with hypothetical proteins of *P. shigelloides*. The VC/PS1 cluster consisted of 19 nodes and 86 edges, with 5 virulence genes and 14 hypothetical proteins (**Figure 7**). The functional characteristics of this cluster were based on gene ontology, KEGG pathways and subcellular localization. VC/PS2 cluster contained network of 41 nodes and 657 edges of 2 hypothetical proteins and 39 virulence genes **(Figure 7)**. This cluster associated with several biological functions, molecular function, cellular component, KEGG pathways and subcellular localization. The smallest cluster of this network, VC/PS3 had 4 nodes and 3 edges which contain 3 virulence genes and one hypothetical protein **(Figure 7)**. The function of this cluster was based on local networking with Type II protein secretion system and Pilus organization.

**Figure 7:**
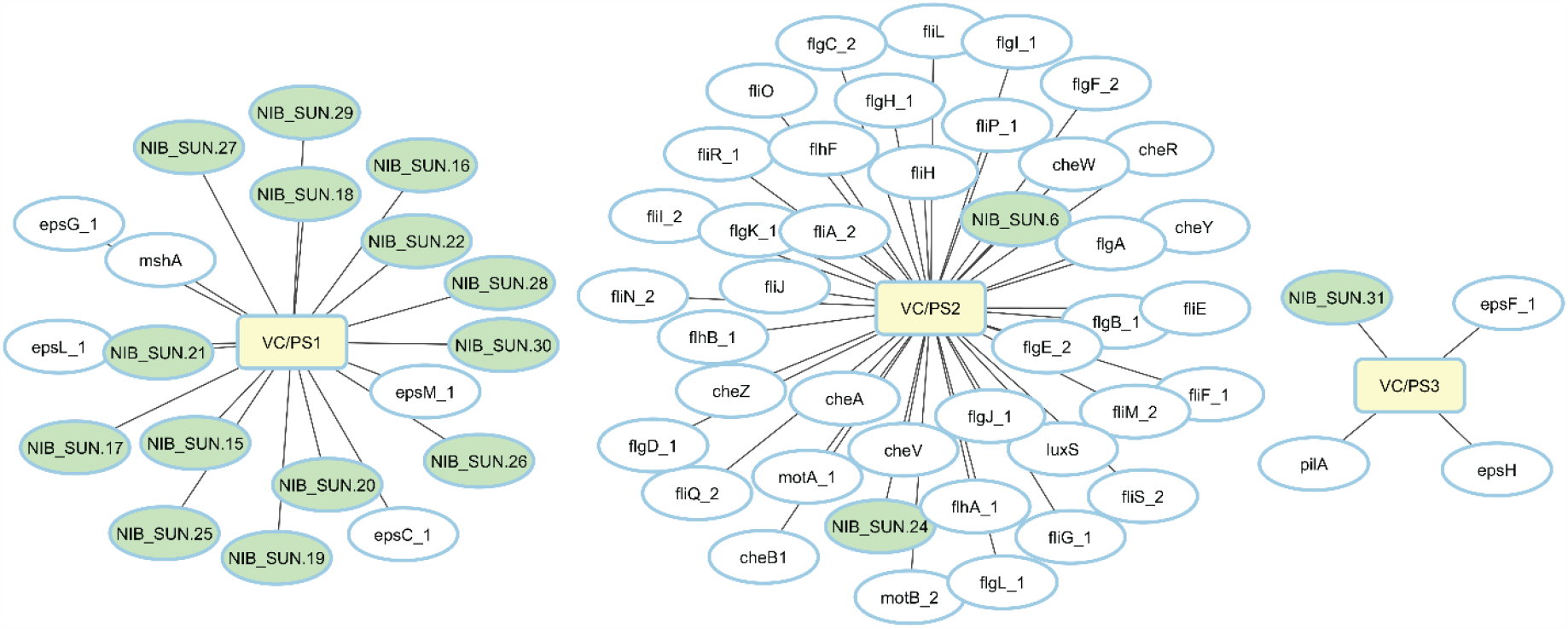
Microbial interaction of *Vibrio cholerae* virulence genes with *Plesiomonas shigelloides*. VC/PS1, VC/PS2 and VC/PS3 are cluster name; Green nodes: Hypothetical proteins of *P. shigelloides*, and White nodes: virulence genes of *Vibrio cholerae*.

The virulence genes of Vibrio parahaemolyticus were shown to form a network clusters (VP/PS) upon interaction with hypothetical proteins of *Plesiomonas shigelloides*. VP/PS cluster was comprised of a total of 9 nodes and 16 edges, including 4 virulence genes and 5 hypothetical proteins, as seen in the **Figure 8A**. This cluster is connected with biological processes, cellular components, KEGG pathways, and several local network, involving flagellar assembly and taxis.

**Figure 8:**
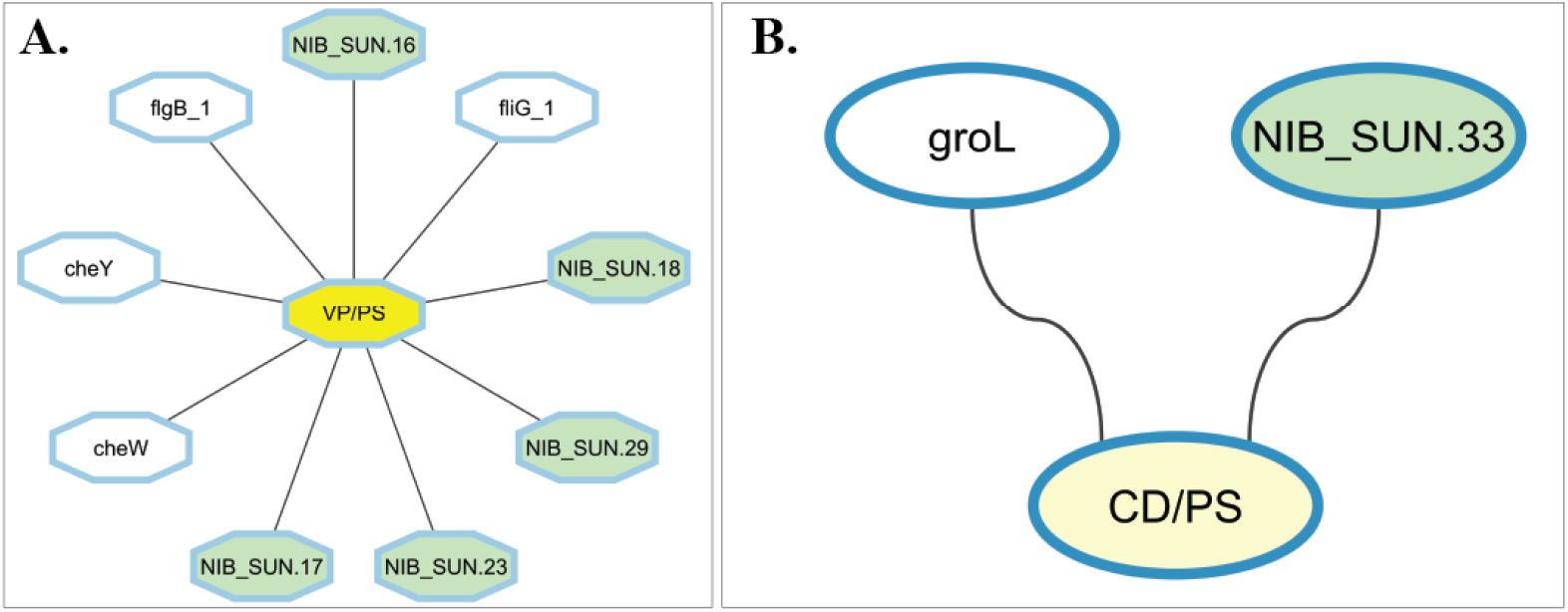
Microbial interaction of (A) *Vibrio parahaemolyticus* and (B) *Clostridium difficile* virulence genes with *Plesiomonas shigelloides*. VP/PS and CD/PS are cluster name; Green nodes: Hypothetical proteins of *P. shigelloides*, and White nodes: virulence genes of (A) *Vibrio parahaemolyticus* and (B) *Clostridium difficile*.

Among 10 common virulence genes, only one virulence gene (groL: Prevents misfolding and promotes the refolding and proper assembly of unfolded polypeptides generated under stress conditions) of *C. difficile* have showed interaction network with one hypothetical protein (NIB_SUN.33) based on their co-expression (**Figure 8B**).

## 4. Discussion

Diarrhea is the most prevalent disease in the world and the second leading cause of mortality among infants under the age of five. In contrast, there are no effective medications or treatments available to eliminate the cause of diarrhea. Researchers are still attempting to find out the most effective drugs and therapeutic targets against diarrheal pathogens. Recent advances open new windows to focus on this purpose. Juan *et al* [15] suggested that multipathogenic infections or coinfections including *Plesiomonas shigelloides* may play a role in the pathogenesis of infectious diarrhea. To address this issue, the coinfection pattern between *P. shigelloides* and other diarrheal pathogens was investigated in this study. Consequently, the identification of hypothetical proteins that possess the potential to serve as drug targets for effectively combating the pathogenesis of *P. shigelloides* is of paramount importance. This research aims to investigate the coinfection pattern as a basis for identifying these proteins.

In this connection, we first collected clinical samples from children and isolated the bacteria using MacConkey and Salmonella-Shigella agar medium (**Figure 1**). Then we identify the bacterium using several biochemical tests and the 16S rRNA sequencing method. Our sequence was 100% similar to NR_044827.1 *Plesiomonas shigelloides* strain. After identifying *P. shigelloides*, we extracted the bacterial DNA and conducted it to the next-generation sequencing platform. The sequencing method was done by using the Illumina MiSeq system and the quality of the sequencing data was good enough. We assembled and annotated the raw sequencing data and found a total genome size of 3,617,790 base pairs having 51.92% GC contents which is almost similar to Edwards *et al* findings [31]. Our annotated results contain 3099 coding DNA sequences (CDSs), that is similar to Alexander *et al* result [32]. We have found 899 hypothetical proteins and 33 hypothetical proteins were selected for the study based on STRING mapping. As our study aims to have interactions with other diarrheal pathogens, we retrieved 100 WGS sequencing data of 10 distinct bacteria, and these bacteria were isolated from stool samples of diarrheal patients in the Asian region. After assembling these WGS sequences, common virulence genes were selected from each bacterial species. Among these, 33 hypothetical proteins were established in several networks and clusters with the virulence genes of 10 distinct diarrheal pathogens.

In interaction network with *Shigella spp*., NIB_SUN.1, NIB_SUN.2 and NIB_SUN.3 established a cluster with three key virulence genes fur, phoP, and rpoS which are responsible for transcription regulation and DNA binding (**Figure 4A)**. We also observed the same cluster in the interactions with *Escherichia coli* (**Figure 5**). These all interactions were based on gene neighborhood and co-expression patterns. These findings provide valuable insights into potential cross-species interactions in *Shigella* sp. and *E. coli* pathogenicity.

In the case of non-typhoidal *Salmonella spp*. 5 hypothetical proteins established two distinct network clusters with 8 virulence genes. The NTS/PS1 cluster is comprised of five nodes and eight edges, which represent the presence of various virulence genes (csgD, csgE, csgF, csgG) and a hypothetical protein (NIB_SUN.22) (**Figure 4B**). This cluster is associated with the biological process of pilus assembly and protein transport, molecular function of identical protein binding and subcellular localization of cell envelope. On the other hand, the NTS/PS2 cluster network consists of 8 nodes and 7 edges (**Figure 4B**). This network encompasses 4 virulence genes, namely fur, phoP, phoQ, and rpoS, as well as 4 hypothetical proteins, specifically NIB_SUN.1, NIB_SUN.2, NIB_SUN.3, and NIB_SUN.32. The inclusion of these genes and proteins in the cluster is based on various factors such as gene neighbourhood, gene fusions, gene co-occurrence, co-expression, and protein homology. So, during diarrheal disease, they might have the same function and support each other.

The most interacting network was built between the virulence genes of *V. cholerae* and hypothetical proteins of *P. shigelloides*. In this network, 47 virulence genes and 19 hypothetical proteins were involved, and they established 3 different network clusters (**Figure 7**). The first cluster, VC/PS1 is involved in gene ontology and has role in bacterial secretion system pathways. VC/PS2 cluster was the most important cluster in this network and this cluster had 41 nodes and 657 edges. Two hypothetical proteins and 39 virulence genes established this cluster. It is associated with several biological functions, especially chemotaxis, protein secretion, protein transport, response to stimulus and cellular process. This cluster also has a molecular function on cytoskeletal motor activity and is involved in KEGG pathways for flagellar assembly, bacterial chemotaxis, and two-component system. In the third cluster, VC/PS3 was associated with one hypothetical protein and three virulence genes, and the function of this cluster was based on local networking with Type II protein secretion system and Pilus organization.

The second most interacting network was formed among virulence genes of C. jejuni and hypothetical proteins of *P. shigelloides*, forming three distinct clusters (CJ/PS1, CJ/PS2, CJ/PS3) (**Figure 6**). The CJ/PS1 cluster consists of 9 virulence genes and 4 hypothetical proteins and this cluster is responsible for the carbohydrate metabolic process and lipopolysaccharide biosynthesis. The most important cluster of this network is CJ/PS2, which consists of 10 hypothetical proteins and 32 virulence genes. This cluster involved several biological processes including chemotaxis, response to external stimulus, and protein transport. It also has molecular function and an important role in KEGG pathways for flagellar assembly and bacterial chemotaxis.

In the context of *Vibrio parahaemolyticus*, 4 virulence genes and 5 hypothetical proteins formed a cluster VP/PS which have several functions (**Figure 8A**). This cluster was connected with KEGG pathways for bacterial chemotaxis and also involved with biological process, cellular component, and some local network clusters.

Only one virulence gene (groL) of *C. difficile* interacted with one hypothetical protein (NIB_SUN.33) of *P. shigelloides* (**Figure 8B**). This cluster is formed based on co-expression and groL is responsible for the molecular function of ATP-dependent protein folding chaperone.

After analysis of all clusters and networks, we can say that during bacterial diarrhea, these 33 hypothetical proteins of *P. shigelloides* might have a pathogenic role with these diarrhea-causing bacteria. Hypothetical proteins may interrupt or stimulate the virulence gene or simultaneously target host cells and that’s why diseases are more severe than single infection of diarrheal pathogens.

Though this study has had some significant findings, we assume that the present study still needs to involve some experimental validation such as Y2H technique, microbial cell culture, western blotting and transcriptomics to justify these findings. Besides sample size, biochemical test and limitations of the proper bioinformatics pipeline were also a hindrance to the substantial improvement of this study.

## 5. Conclusion

The present study has integrated in vitro and in silico approach to characterize the hypothetical proteins of the *Plesiomonas shigelloides*. Further, microbial interactions were scrutinized through the systems biology approach. The findings of this study suggested that 33 hypothetical proteins, which are present in the WGS, could interact with the other diarrhea causing bacteria in terms of their pathway, localization, biological and molecular function connection. Therefore, these 33 hypothetical proteins might be a potential drug target against diarrheal pathogens when it comes to broad-spectrum drug effect. In addition to these findings, the present study has laid the foundation to explore the microbial interactions between *P. shigelloides* and other diarrhea causing bacteria for the researchers.

## 7. Author Contribution

Mohammad Uzzal Hossain, A.B.Z Naimur Rahman, Md. Shahadat Hossain, Shajib Dey: Conceptualization; data retrieval; sequence analysis; laboratory experiments; writing – original draft; Zeshan Mahmud Chowdhury, Arittra Bhattacharjee: writing – review and editing; Ishtiaque Ahammad: manuscript preparations; Khandokar Fahmida Sultana, Abu Hashem, Keshob Chandra Das: supervision; validation; Chaman Ara Keya: supervision; validation; Md. Salimullah: project administration; validation; supervision.

