## Supplementary file for "Unraveling Coinfection Dynamics into 100 Whole Genome of Diarrheal Pathogens: A Genome-to-Systems Biology Approach with *Plesiomonas shigelloides*"

**Supplementary Figure 1:** **Biochemical test of *P. shigelloides*.**

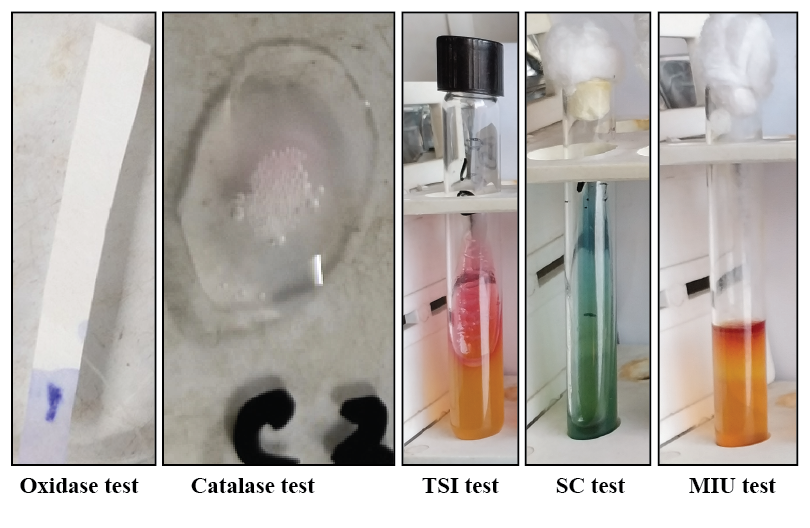

**Supplementary Figure 2:** **Assembly quality of the *Plesiomonas shigelloides***

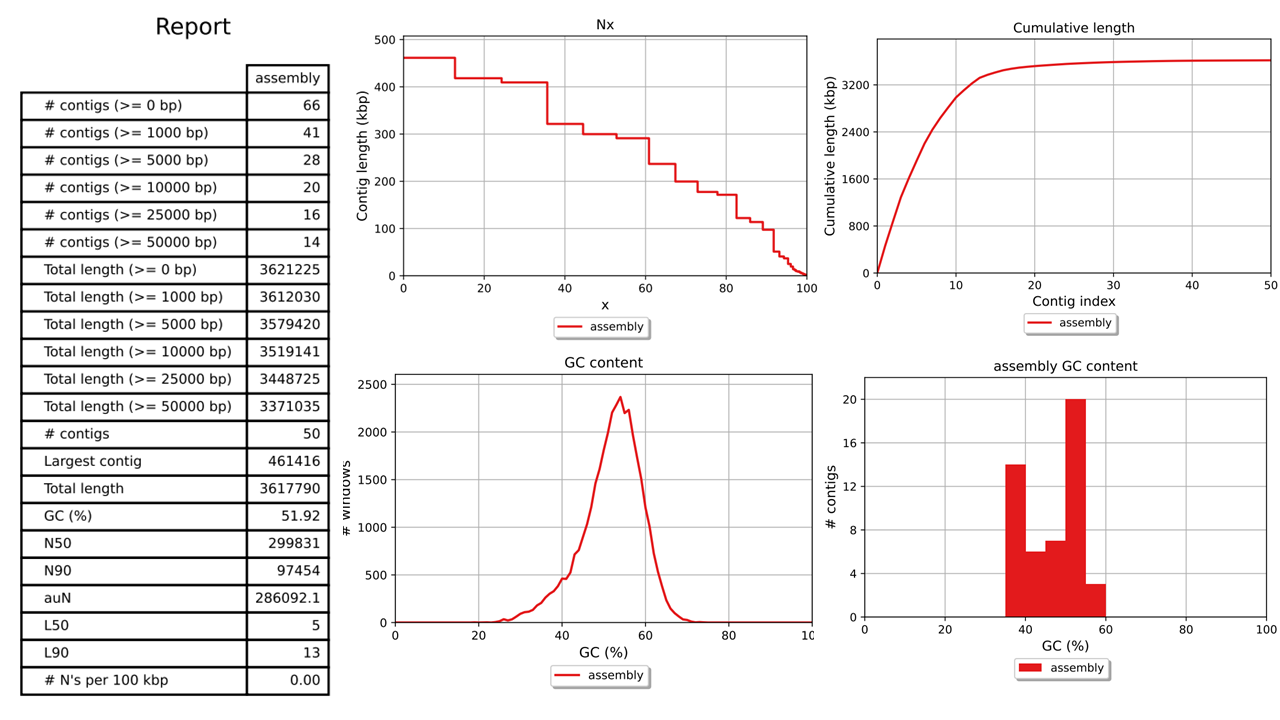

**Supplementary Table 1: Sample metadata information.**

| Sample ID | Age | Sex | Country |
| --- | --- | --- | --- |
| H-1 | 0Y 4M | Male | Bangladesh |
| H-2 | 0Y 10M | Male | Bangladesh |
| H-3 | 1Y 3M | Male | Bangladesh |
| H-4 | 0Y 6M | Male | Bangladesh |
| H-5 | 0Y 4M | Male | Bangladesh |
| H-6 | 0Y 1M | Female | Bangladesh |
| H-7 | 0Y 1M | Female | Bangladesh |
| H-8 | 0Y 10M | Female | Bangladesh |
| H-9 | 4Y 6M | Male | Bangladesh |
| H-10 | 2Y 6M | Female | Bangladesh |
| H-11 | 8Y 6M | Female | Bangladesh |

**Supplementary Table 2: SRA dataset information of 100 whole genome sequences.**

| **No** | **Bacteria** | **Country Name** | **SRA ID** | **Isolation Source** |
| --- | --- | --- | --- | --- |
| 1 | *Campylobacter jejuni* | Japan | DRR301692 | Human, stool |
| 2 | *Campylobacter jejuni* | Japan | DRR301693 | Human, stool |
| 3 | *Campylobacter jejuni* | Japan | DRR301694 | Human, stool |
| 4 | *Campylobacter jejuni* | China | SRR11249039 | Human, stool |
| 5 | *Campylobacter jejuni* | China | SRR11249040 | Human, stool |
| 6 | *Campylobacter jejuni* | Bangladesh | SRR11268219 | Human, stool |
| 7 | *Campylobacter jejuni* | Bangladesh | SRR11268220 | Human, stool |
| 8 | *Campylobacter jejuni* | Bangladesh | SRR11268221 | Human, stool |
| 9 | *Campylobacter jejuni* | India | SRR11268152 | Human, stool |
| 10 | *Campylobacter jejuni* | India | SRR11268153 | Human, stool |
| 11 | Enterotoxigenic *E. coli* | China | SRR14868194 | Human, stool |
| 12 | Enterotoxigenic *E. coli* | China | SRR14868195 | Human, stool |
| 13 | Enterotoxigenic *E. coli* | China | SRR14868196 | Human, stool |
| 14 | Enterotoxigenic *E. coli* | China | SRR14868197 | Human, stool |
| 15 | Enterotoxigenic *E. coli* | Bangladesh | SRR8634233 | Human, stool |
| 16 | Enterotoxigenic *E. coli* | Bangladesh | SRR8634248 | Human, stool |
| 17 | Enterotoxigenic *E. coli* | Pakistan | SRR8634234 | Human, stool |
| 18 | Enterotoxigenic *E. coli* | Pakistan | SRR8634235 | Human, stool |
| 19 | Enterotoxigenic *E. coli* | India | SRR8634249 | Human, stool |
| 20 | Enterotoxigenic *E. coli* | India | SRR8634250 | Human, stool |
| 21 | Enteroinvasive *E. coli* | Bangladesh | SRR10572924 | Human, stool |
| 22 | Enteroinvasive *E. coli* | Japan | SRR13615174 | Human, stool |
| 23 | Enteroinvasive *E. coli* | Thailand | SRR2994334 | Human, stool |
| 24 | Enteroinvasive *E. coli* | Thailand | SRR2994360 | Human, stool |
| 25 | Enteroinvasive *E. coli* | Thailand | SRR2994368 | Human, stool |
| 26 | Enteroinvasive *E. coli* | Thailand | SRR2994369 | Human, stool |
| 27 | Enteroinvasive *E. coli* | Thailand | SRR13614963 | Human, stool |
| 28 | Enteroinvasive *E. coli* | Thailand | SRR13615153 | Human, stool |
| 29 | Enteroinvasive *E. coli* | India | SRR11606889 | Human, stool |
| 30 | Enteroinvasive *E. coli* | India | SRR11606891 | Human, stool |
| 31 | *Salmonella* Typhimurium | China | SRR10321788 | Human, stool |
| 32 | *Salmonella* Typhimurium | China | SRR10321789 | Human, stool |
| 33 | *Salmonella* Typhimurium | China | SRR10321790 | Human, stool |
| 34 | *Salmonella* Typhimurium | Hong Kong | SRR13527909 | Human, stool |
| 35 | *Salmonella* Typhimurium | Lebanon | SRR8564894 | Human, stool |
| 36 | *Salmonella* Typhimurium | Lebanon | SRR10018689 | Human, stool |
| 37 | *Salmonella* Typhimurium | Lebanon | SRR8544183 | Human, stool |
| 38 | *Salmonella* Typhimurium | Taiwan | SRR12831475 | Human, stool |
| 39 | *Salmonella* Typhimurium | Taiwan | SRR8189410 | Human, stool |
| 40 | *Salmonella* Typhimurium | Taiwan | SRR15827378 | Human, stool |
| 41 | *Salmonella* Enteritidis | Bangladesh | SRR19174821 | Human, stool |
| 42 | *Salmonella* Enteritidis | Bangladesh | SRR19174822 | Human, stool |
| 43 | *Salmonella* Enteritidis | India | ERR6719637 | Human, stool |
| 44 | *Salmonella* Enteritidis | India | ERR6719640 | Human, stool |
| 45 | *Salmonella* Enteritidis | India | ERR6719644 | Human, stool |
| 46 | *Salmonella* Enteritidis | Cambodia | SRR10454708 | Human, stool |
| 47 | *Salmonella* Enteritidis | China | SRR15962854 | Human, stool |
| 48 | *Salmonella* Enteritidis | China | SRR15962859 | Human, stool |
| 49 | *Salmonella* Enteritidis | Thailand | SRR1532573 | Human, stool |
| 50 | *Salmonella* Enteritidis | Thailand | SRR1532580 | Human, stool |
| 51 | *Shigella dysenteriae* | Pakistan | ERR6005829 | Human, stool |
| 52 | *Shigella dysenteriae* | Pakistan | ERR6005830 | Human, stool |
| 53 | *Shigella dysenteriae* | Pakistan | ERR6005831 | Human, stool |
| 54 | *Shigella dysenteriae* | Bangladesh | ERR6005841 | Human, stool |
| 55 | *Shigella dysenteriae* | Bangladesh | ERR6005842 | Human, stool |
| 56 | *Shigella dysenteriae* | Bangladesh | ERR6005843 | Human, stool |
| 57 | *Shigella dysenteriae* | Bangladesh | ERR6005844 | Human, stool |
| 58 | *Shigella dysenteriae* | India | ERR6005869 | Human, stool |
| 59 | *Shigella dysenteriae* | India | ERR6005870 | Human, stool |
| 60 | *Shigella dysenteriae* | India | ERR6006254 | Human, stool |
| 61 | *Shigella flexneri* | Bangladesh | ERR6004552 | Human, stool |
| 62 | *Shigella flexneri* | Bangladesh | ERR6004553 | Human, stool |
| 63 | *Shigella flexneri* | Bangladesh | ERR6004554 | Human, stool |
| 64 | *Shigella flexneri* | Bangladesh | ERR6004555 | Human, stool |
| 65 | *Shigella flexneri* | Pakistan | ERR6004631 | Human, stool |
| 66 | *Shigella flexneri* | Pakistan | ERR6004632 | Human, stool |
| 67 | *Shigella flexneri* | Pakistan | ERR6004633 | Human, stool |
| 68 | *Shigella flexneri* | India | ERR6004615 | Human, stool |
| 69 | *Shigella flexneri* | India | ERR6004616 | Human, stool |
| 70 | *Shigella flexneri* | India | ERR6004617 | Human, stool |
| 71 | *Vibrio Cholerae* | Bangladesh | DRR394994 | Human, stool |
| 72 | *Vibrio Cholerae* | Bangladesh | DRR394995 | Human, stool |
| 73 | *Vibrio Cholerae* | Bangladesh | DRR394996 | Human, stool |
| 74 | *Vibrio Cholerae* | Bangladesh | DRR394997 | Human, stool |
| 75 | *Vibrio Cholerae* | Thailand | ERR1485195 | Human, stool |
| 76 | *Vibrio Cholerae* | Thailand | ERR1485196 | Human, stool |
| 77 | *Vibrio Cholerae* | Thailand | ERR1485197 | Human, stool |
| 78 | *Vibrio Cholerae* | India | ERR5032476 | Human, stool |
| 79 | *Vibrio Cholerae* | Taiwan | SRR7791597 | Human, stool |
| 80 | *Vibrio Cholerae* | Taiwan | SRR7791599 | Human, stool |
| 81 | *Vibrio Parahemolyticus* | Malaysia | SRR24070348 | Human, stool |
| 82 | *Vibrio Parahemolyticus* | Bangladesh | SRR1106412 | Human, stool |
| 83 | *Vibrio Parahemolyticus* | Bangladesh | SRR1106435 | Human, stool |
| 84 | *Vibrio Parahemolyticus* | Bangladesh | SRR1106438 | Human, stool |
| 85 | *Vibrio Parahemolyticus* | Thailand | SRR1060690 | Human, stool |
| 86 | *Vibrio Parahemolyticus* | Thailand | ERR5319171 | Human, stool |
| 87 | *Vibrio Parahemolyticus* | India | ERR5319173 | Human, stool |
| 88 | *Vibrio Parahemolyticus* | India | ERR5319186 | Human, stool |
| 89 | *Vibrio Parahemolyticus* | Hong Kong | ERR5319187 | Human, stool |
| 90 | *Vibrio Parahemolyticus* | Hong Kong | ERR5319188 | Human, stool |
| 91 | *Clostridium difficile* | South Korea | SRR18672521 | Human, stool |
| 92 | *Clostridium difficile* | South Korea | SRR18672522 | Human, stool |
| 93 | *Clostridium difficile* | South Korea | SRR18672523 | Human, stool |
| 94 | *Clostridium difficile* | South Korea | SRR18672524 | Human, stool |
| 95 | *Clostridium difficile* | South Korea | SRR18672525 | Human, stool |
| 96 | *Clostridium difficile* | China | SRR7300068 | Human, stool |
| 97 | *Clostridium difficile* | China | SRR7300069 | Human, stool |
| 98 | *Clostridium difficile* | China | SRR7300070 | Human, stool |
| 99 | *Clostridium difficile* | China | SRR7300071 | Human, stool |
| 100 | *Clostridium difficile* | China | SRR7300072 | Human, stool |

**Supplementary Table 3: List of hypothetical proteins of *Plesiomonas shigelloides***

| **NIB ID** | **Amino acid sequences** |
| --- | --- |
| NIB_SUN.1 | MPAFLFRAWIFRWALNLWPPFFGAGIRIRRLDRDFRCCEVTLAFRWWNKNINRSQFGGSLFAMTDPIYALMLIGRLGPDYMVWDKAADIDFIRPGWGKLTASFEISEGRYEEILAATANGEKCLPQFTVAIKDAQGEIVARVNRTLYVRRKRKTEEVVAPLDAHRPHFATGDKPHEKNKAG |
| NIB_SUN.2 | MSTELTWDQTLRFRLLEIVLQWEGRLTTNHLCTAFNIGRQQASRDINRYKNDFAPQGLEYDHSLKGYRPTAEFKPQFSQGKVDEYLTLLHREKQHAVQSQQQALLTLPLLDLRYGHIEMLDVPTRHVDPVIVRGLVQAARGQLRVDVNYVSLSSEEESGRNIVPHTLVYDGIRWHVRAYCEKKGEYLDFVMSRFRGEPDLLDASEHGREHDVEWNTLVTAEIIPNPHLSAGQQAIIAADYAMDDGHLLLTQRMPLMHYALERMQVSFDGEHHQQPAHAPIILKNRAELIAQGCTFKLKSADLQPSIAVASTAYA |
| NIB_SUN.3 | MYSRHAQQRMQQRGISSQAVEWLLDFGHADYHRGREIFFWSRKSLARLAQQTSISHQQCQRLRRHYLVLDDGEIVTVGHRTTHFKRDRH |
| NIB_SUN.4 | MLIEIRALDTRLRELFAPDADPDPDEILQLMGQRQQLLQRLIPTLSVENKQQLLLETQSLLCLAQTAKQECGDKLAVRKRGQRGINAYRQVSTQ |
| NIB_SUN.5 | MRAGLISLLLLAAGFAQAEVQVAEPYVRATVPGAQMSAAFATLFNPEDDERHLVAVSSPVAKQVELHDMVVSPDGVMNMQKVEQITIPAKGTLMLKPGSLHIMLLQLTQPLPAGTEVPLTLTFDNGQQVQIQATAREVKPAGHHHPKS |
| NIB_SUN.6 | MSGLSERLQRLSQAMEQAVRSGDFTLLAELDQKLLAGLSRLNGAEIPADAQPVLKQLASRYPQWIAAARSEQAQVRAQICQQPSQQQGLQAYIQNS |
| NIB_SUN.7 | MNRIVMALLVALGLAAGGWLAGYGVEKSRAPVRTVTVKGLAEQDVKADLALWPIRFVRAGNDLAQVRDEIQRDDTAVTAFLQKNGLADAVSFRGMDVTDRNAQQYGDGNVPFRFIITRTIMLRTQDVAKIDEAYKQSSQLVDSGVILSNEGVGSGNPVYLYNGLTTLKPAMIAEATKNAREAAQQFAHDSQSRLGRILVADQGVFQILPRDNAPMLQEDQQIDKTIRVVTTIKYQLVD |
| NIB_SUN.8 | MVNQVNSTRLGTQRTSDLNHSPVVSQQKAGLAVSASASAAVDTRTPAEEIERLRHGQRGSLLGVTAKAHYQLAKTQTALRALHSAEDILQQSKQLAQRSLKQTEGRRQRLTQELASLQRQLRQQVHQTYDEKPLFNADFSVRHLNRNQQQRFSIRGLRLQGERSQPEVLKFAFDGGKPRQVKVQIEPEDDMVTVARKLSRGLSSRGIRAEVDKWGDLEFSAAKGNWQEVEEGFWLTGNGQIFPAGNPVKVQPEQRQPQMVDAEKLSFRKTQDTRQSLADIERLMQQIRSSLQELTRYQQQIERQLQGRYARMPETETAGISDRIVNKPLSSLLAQAGISRQTVVALLTR |
| NIB_SUN.9 | MPLDNYFRDELAYLRLQGREFADAHPELTRFLSEQTTDPDVERLLEGFAFLTGSLRAKIE  DEFPELTHGLLGMLWPNYLRPVPSMTVMQFSVIPGAIVQPAPVARGCRLDSLPVDGVTCHFQTCHDAWIYPADIHDIRAQSGNDLSTITTEIHLHSPLTLAELQLDKLRFYLGGDSYTANELYYWLSAQLSYIEFEVNGQRFRQSASVLKALGFERDDALLPYPSNVYSGYRILQEYFCFPESFLFFELSGAGADWPKQPLAVTEFKLHFCFERPLPSALKIRPDSLQLNCVPAINLFQHDSEPINLHGRQTEYPLKASYRYADSFEIFSVDKVDSWAEGGIGRVRGATRQYHPFESFQHQIERAKGRQALYYRIRVREAVSGDGFEHALSFVRGDETAMLGLDESVSVSLTCTNRGKAARLPVGAICVPTGSSPSFATFKNLLRPTRPLRPALDGSLHWTLISNLSLNYVSLLRRDALVQILRTYDFPALHDKQAEQASRKRLAGIEQIETVPVDRLVRGMPVRGLKSVLSIRQSAFGSEGELYLFSTVLAHFFSLYASVNAFHLLEVVNLDNQERYRWPVQIGQHSMM |
| NIB_SUN.10 | MLRTLWCAVWLLALTGCTTLGKMADVAMNPDIQVGSNADQPSTVGLSLLAEPDVNPNESGEAAPIEFQVLLLSEDSRLLATDYDQVTEDVAKVLGKNYLDHQDYTLLPGQFKYLPPVKLEEEVRYIGVIARYAEPDKAEWRKVIKVKNTGRHYQILVHLRQDEVELQKEEE |
| NIB_SUN.11 | MSVVEEQVKAGAQASSGSLLDEIMAQARITPVDEGYSVAKQGIAALVANILDSGNTAEPVNKALVDSMIVELDKKLSKQMDVILHAQPLKELESSWRSLKLLVDRTDFRENIKIQVLHATKEELLDDFEFAPEITQSGFYKHVYSSGYGQFGGQPVGAVIGDYAFSQSSPDIKLMQYVSAVGSMAHAPFISSVAPTFFGVDSFTDLPSIKDLKSVFEGPAYTKWRSLRESEDARYLGLTAPRFLARLPYDPVENPIKTFNYKEDISADHDHYLWGNTAYLMGSALNDSFAKYRWCPNIIGPQSGGAVTDLPVHVYEAMGQLQAKIPTEVLITDRREYEMAEEGFITLTMRKDSDNAAFFSANSVQKPKVFPNTKEGKEAETNYKLGTQLPYMFIINRLAHYIKVLQREQIGSWKERQDLERELNGWIKQYVADQENPPADVRSRRPLRAAQIKVLDVEGEPGWYQVAMSVRPHFKYMGASFELSLVGRLDKE |
| NIB_SUN.12 | MSSRNRVIWREGLFIKPQHFQQQQRHTDHMLHARLSALSDYFYGLQSLSINEDYLGFGRIALVSSSGVMPDGTVFSIPVDDVLPPPLEVTDVSVANQKVYLALPLSVSGVSEVGESGQAVATRLQSHRHDLRDLHSDGGDITSLEVGKVSLRLMLEREDRSAYAAIAIARILDKRPDGGLVLDPNFMPCSISVAAIPTLKRFLGESAGLVAERARSLAQRLAAPGQQGVADVAEFMMLQLLNRAQPQLSHLARLGTLHPERLHEALVQLCGELMTFTDESRLPPEFPAYRHDDQQVSFEPVILALRQSLSTVLSPRAVSIQLRKHPYGIMVAMVGDSDLMASAEFVLAVRARMPQEQLRKQLLQQTKIASSEKIRELISLQLPGIPLLPLPVAPRQLPYHAGYSYFQLDRQSPAWQMLAQSNTLAFHIAGDFPELDMQFWAIRSQG |
| NIB_SUN.13 | MTIDIIQHEQLDDLLYDHAAQLDLDSDYWFRLRGHSINPMIDAVTPILGMVLRVRQLSSHSQVGELYQRVVTEIQAIEQELMAHGYENGVVLSFRYILCTFIDEAVMSREWGSQSEWSEHSLLTRFHNETWGGEKVFVLLGKLQEDPTRYRDVLEFIYLCLCLGFEGRYRVMSHGRDEFERVVRQLHRQLYPDGRDAPTVFHLDLGQQASRYQLKRQVSLRSLCIGAVLLLTGIFVLYYTRLDNQTQDVLRQLSDLLK" |
| NIB_SUN.14 | MRLNQAVKMVAGVVLALALVGCGARGDTPAQKRNAIHQMRQDTLQKLYREQPKARTAIRKAQGYAVFSNKSNKLGFIGIGHGFGVVRDNRTGKNTYMRMMSVGAGVGLGIQDIRFVAIFHDRKTMQNFIQHGWDASAGADAAAKWKEKGDAGNQTVAADFNGVTIYQLTEHGLALQAMVQGYKYWPDDELN |
| NIB_SUN.15 | MRKWLGVIGLCGLLLGCASSGSRQTLTAVGYAPIATQSGKTLQERQIQAMRASRLDAYRDMAEQVYGLRINTDSQVAADGALRDAGIRSQAAGVIRGAEVVRSYPVGDNYVTELRLDMSQVKQLPPLPAPAKPAAPAPQPVILVPARGATY |
| NIB_SUN.16 | MHNGLDNGATITARMGITRWSNRLTRPRRLAFRMLACCLGMTLSACVAAMDSDATAPMTPPDSITVEHDAIFANRFRLDRTVQWLTLLISRSDNQEQLVLVQPDGSKLNARMDSPDVHWWRSPTLEIVTINRPMQGPWQVSAPEGTVSTFIGVPDMQLSADSMPQPLYQHEQLKLTARVLVNNQPLLRADILQHLRLHAEFVPVNANHALQAGQDTVRAGRPAVMAGELRDDGQHLDAKAGDGIFTMQLPVNPAPGEYVLTINTGDGIYLPPFREQVTVHPMPLQAQFLQGRTASQNHRLDVSSDATRIVPGSLLVRVDEKIPGGLNAEYQVQAPATGNSVSLEIPQRDFIGNYSLTAWIYAQQVGDRPLSLLLPEQRFMVVTSQQQTQQLLNQLAQQKQQQALSEQAQAELASVTQAARNQGSHRLQMVLLGTVLVVGIFGAAIWGGMALRRAHQRRQALQVLRHQQASSQKVGQRDLSSSR |
| NIB_SUN.17 | MEQAQFFSVHQHLSVNIIPLRADETLPDDARFDEEIPLPFRIAAETAHLDISSIRSLRNIGSAADDLMGFLRIQQQKIDLVVNYILAQEDQPHLRHLTHSFSAGGFSYFADTPPASGSIMRIKLFLSQPAAAVYAYGEVSEVSIEDGRPLVQIRFIRLRDKDQDLLIRAALHEQQQQLKQRAELRVKD |
| NIB_SUN.18 | MNLNRDCPRQWLLPVITVPVGDVMSNAINVPFHPVQVFTVTGDKITENVTGTQNVKHMKTLLPLAALSGLLLTGCAGNQPSADQPYVMDKNQVQVLPRHSLDFFVDAMARQLVDSNKTLAASGPIAVASFVDLQQMDQTNWLGNLMADSFIYQLQQRGMTVLDYKTTGQIRVTPQGDFALSRNWRDLQKAYQVNSVLTGTMLRQGSGIQINARIIRFSDRVVIATAQGFLPADRLGRSIGNPNLVQMRNGMIVRGDNRFTSPSIVAP |
| NIB_SUN.19 | MIGLMVAVAVLSVVCAGLAWSLISLWRQHKTILRRVEASEVLLKAARRQFDTTGKSIEELRGGFIGLGNELHQLVDKQEQFSVKLNEVTYIDPEHRLYSRATKLAEMGAGLDELMQECELPKAEAELLLNLQRQMKNRRT |
| NIB_SUN.20 | MIYLIAFGIFGIMILLMMIGQLLTKRQIQGSCGGLSQAGIERVCDCEETCAEHRLYQISEPSAAPSAPEAAASKTAQQ |
| NIB_SUN.21 | MTSAEQAAGQESSASANEHTDNSRLISARQRAIDLAINRHIDGKIYPQQPTKLEQRLQRRLQQEQLQRQENLEKIMQIAASLCQDTLATESDPDWISSFLQLAQETRGAPMQRLWGRILAQEIVQPGSFSIKALRTLRDMTHREAMLFQRACAVACSFGADLSAKRIITGIRPGKNGLKSLFRAPAPEQLNLGSFKLPFSALLLLQELGLLLGGELESGSLLPEQNYSLRFASTDLTLTTHDTGHRLTYYRFSPVGEELSQLLIGTSLPEYIQCLSSLLGAHLTVTPRSANVSV |
| NIB_SUN.22 | MTLTLRPRATCLATVLMLASSLYTSNLYAAWYQTTVSIPTQAGSESQLRSLALSDASGQL LGYAGVSMNYQQPLVASLLSGDKDGQTIRYAAGSPVHAATVVSSATQNGRFTLTLRADITPDRSLCRRQGYRKPLAVYRFALNDLTQASLGAIFNIDSDYPRVLERQLRQDLQGLQVRSYQNEHLPADASANWLAAAGEQQQAQYLLTGQLIDLSPAPVTGLNTFFRVSPMKRHFAVNISVFDSYSGEQVFERLYRTEAEWPFERTERVDTSSANFWQSAYGYAIRLQTDQIAQDLDTALGCSATQARITEINGNRLKINIGSRNGVRKGDRVKLLHTTEFVDQSGNVAEMFTPSGMSLTVSQVYATYAIINIGSLNTASTIQVGDLVTKS |
| NIB_SUN.23 | MLMVRDTLRSITLWKTGIALLCLSLSPLAQALTKLDISIDKEPVEQGQPFTLTITADDLLPNEALDLSPLFGQSFVVGNTEYNQIPADKKQPAATRWKTTLMQDQTGNYQIPALTIDNIASQPLAFNVVPAKVEEKKEEVPAEPAKPVITLDQAVLNTRLSKPVAYPDEAVYLDIDLELASELLDGVLSTPELRNGTLRQVGQDRSWSLMRNGNRYQIVSRRYQIMAHEPGNITISNIRFRGTVRGSSQLQPPLNAYAEAMPVTLNVRPLPKDLAANWLPAEKVSLEETWFPQPTAAKVGEPLTRTIRLRARGLGDKPLVMPEIVYPAELSLHAQPVEQRRTVNGETELEIRHILMPKRTANNVMLPEISLTWWNTQTEKAETAVLAAQPLVISSNPALSQPVVPMVKPKENYANLFGYIAIFGGFLLAGGLYWHFSQKRSLRNELSRLSLMMPSEPEPEEKAWLELEQALNSHEPAQITRALSQWARQRWPGRKFTHFAELPSGDAIDRELTRLRRSKDNKSSWNPQTLRQLLHRIRGIRSYNSNSTPAVSTPVTDLRRDDDFPKISLKLDDKD |
| NIB_SUN.24 | MSVAPKERINIKYVPATGGQQAEIELPMTMMVIGNMKGRVEDTPIEERQTVSIDKNNFSS VMKESGLELNFSVQNRLEEGSQDELPVKLQFASMADFAPDSIANQVPELQKLLELREALVALKGPLGNIPAFRNRLQDLLSSDEAREQLLKELDLVAPAE |
| NIB_SUN.25 | MLASTHHYAHALLRRIAQTWLSVSLRGLVTLLLGSVLLLSTPARAEQLQTLGNLEIHYSVFPSTMLTPEIASRNGLTRSNYRAILTVTALDASVAGKPAVDINLSGQAQNLIGNTRQLSF KAVQEGSARYYLAEVPIVNEETYRFTLDIQSKTASGTVRFQQTFYTD |
| NIB_SUN.26 | MAWQQGLVLAAVMVLLPLGLYAGWLLGRLWRQGQLQRQQQAALAAHRRQHMLDSIRTIARAMQSGQCELAEGVLRIGRLLELLVEQDKPELARDFPAMAELYQVVAEMPTHDARKQHKRNQIMKWDLERVRAEERLQPAIEQELPQLLAWCAKYPE |
| NIB_SUN.27 | MIRPARTSRVILPPLGWSMIPKGERYRELLHQTLAPWWGKIFGFHLLKLGALSADLDSSASPIQHHVAVAPEGPQLSVLADPDCLPFTNKAFDAALLAFTLDYASDPHAVLREVDRVLVDDGYLILIDCNPFSLAGIGRCVPLLRRRFPYQCRMFAPFRITDWLHLLNYEVIHVEAMLPFPGRASSRRLAGFCTCHLPVLGSWRLLVARKRTIPLILTGSKRALNNPRMMYPVSACQQVNSPADSRCHPE |
| NIB_SUN.28 | MTQQVWYKQFWPWFLILLPLAAVAASFYTLKLAITHRDSLVVDDYYNKGKAINEDISRLQLAEQLHIQGDLLIVGEQARLTLKKGDLPAFPALKIYFYHPTLQDHDFNLMINADYNGVYRFDLPSAITGKWQVSIEPFDSRWRLQESITLPSSALFSFSRHR |
| NIB_SUN.29 | MSTILAALRRAEQERRRQPTGEPLLPTLPGGPLLSADSAASAALAQRRRHYQRLGLRLLLLLLGASGMWLGLRYGIGPDTRAPQISTSSVSSIEPRKPTIENVHAQPASVSVPTASALPK AEDHLVQPQVLSFPRLETERISAEQLASALSPERESWALPPPIAGIALDDNLITAADILPTPPLPTDPVRRPPAANSDASALPASAVVSRGLADTDTALTPPDKLDLRGVSPELAQAFRAALQDKSSSVTSGAGRAGSAMPVTRPEITAPTSPTASTKKDASAKALPQLRHLPAARRALI PSLRFEVHNYVSAPDKRWVRINGQVYRSGAQLAKGVQLLRIEPEQIVLAVKGQQAALPALQDWPGSRG |
| NIB_SUN.30 | MTVFIATFAVFVLVIAGMAIGYIVQKKSIHGSCGGIASLGLEKVCNCPEPCDKRKAKMAE EARLAKLQENRII |
| NIB_SUN.31 | MKQYGYIASAVCLLILSGCSTSTKNSVAESIAESQVQKINQSLPAKAAGYTLVFAQQQGA EIGLTLVADNQSAARMSPSAVLAGYQQGLCSDPTVRTYLAKGVSYRLQVNNLQGSSLQAAHLTAAQCASQSYARAVHRAGE |
| NIB_SUN.32 | MNRIGRWCAVTLTVLMMTGFVASQTGADPLPELELDDDVQDTALAATQAGLIQPYSLLLQQVERDLNGRIIEVELDEDDGVWEYELKLLGAANNVIKVKYNATTLKILKIKGHDLDSVLRHGVDLSQLDD |
| NIB_SUN.33 | MILFFLVFIIIMTASTWCIFVKAGYPGWAAIIPFYSTIIILKITHRPLWWIFLLIMPIIG FIISFIVFIDLAKAFGKNVYFGIGLALFGYIFFPILGFGKAKYTGKNI |

**Supplementary Table 4: List of virulence genes**

| **Bacteria** | **Virulence genes** |
| --- | --- |
| *Campylobacter jejuni* | flgA, flgS, fliP, flhA, Cj0883c, ciaB, flgR, fliW, cheY, waaC, htrB, Cj1135, waaV, waaF, gmhA, hldE, hldD, gmhB, fliR, porA, Cj1279c, pseB, pseC, flaC, flgG, flgG2, flgH, rpoN, fliS, fliD, flag, flgB, flgC, fliE, Cj0371, fliN, motA, motB, flhB, fliH, fliG, fliF, cheV, cheA, cheW, pflA, fliQ, hddC, gmhA2, hddA, Cj1427c, rfbC, Cj1420c, Cj1419c, Cj1418c, Cj1417c, Cj1416c, cysC, kpsC, kpsS, fliL, pseI, pseA, pseH, pseG, pseF, Cj1442c, kpsF, kpsD, kpsE, kpsT, kpsM, flgI, flgJ, flgM, flgK, cadF, fliK, flgD, flgE, fliY, fliM, fliA, flhF, cdtC, cdtB, cdtA, fliI |
| Enterotoxigenic *E. coli* and Enteroinvasive *E. coli* | rpoS, ompA, fimH, fimG, fimF, fimD, fimC, fimI, fimA, fimE, fimB, espX4, espX5, fur, entA, entB, entE, entC, fepB, entS, fepD, fepG, fepC, fepE, entF, fes, fepA, entD, rcsB, galF, gndA, gspE, espR1, phoP, csgC, csgA, csgB, vgrG, tssI, tssB, tssC, tssF, tssG, tssJ, tssK, tssH, tssM, tssM, espL1, acrB, fdeC, ykgK, ecpR, yagZ, ecpA, yagY, ecpB, yagX, ecpC, yagW, ecpD, yagV, ecpE, espX1 |
| *Salmonella* Typhimurium and *Salmonella* Enteritidis | mgtC, mgtB, misL, lpfA, lpfB, lpfC, lpfD, lpfE, slrP, sopD2, sinH, ratB, ssaJ, sifB, steA, sopD, rpoS, invH, invF, invG, invE, invA, invB, invC, invI, invJ, spaO, spaP, spaQ, spaR, spaS, sicA, sipB, sspB, sipC, sipC, sipD, sipA, sspA, iacP, sicP, sptP, iagB, hilA, hilD, prgH, prgI, prgJ, prgK, orgA, orgA, hilC, sprB, avrA, mig-14, pipB2, ssrB, ssrA, ssaB, ssaB, ssaC, ssaD, ssaE, sseA, sseB, sscA, sseC, sseD, sseE, sscB, sseF, sseG, ssaG, ssaH, ssaI, ssaJ, ssaK, ssaL, ssaM, ssaV, ssaN, ssaO, ssaP, ssaQ, ssaR, ssaS, ssaT, ssaU, basS, basR, steB, sseJ, fur, sseI, sseI, ompA, pipB, sopB, sopB, steC, sseL, rcsB, sspH2, galF, sopA, sodC, sopE2, STM0266, acrB, allB, fimA, fimI, fimC, fimD, fimH, fimF, fimZ, fimY, fimW, fepG, entB, entA, csgG, csgF, csgE, csgD, csgB, csgA, csgC, sifA, phoQ, phoP |
| *Shigella dysenteriae* | rpoS, phoP, gspM, gspL, gspK, gspJ, gspI, gspH, gspG, gspF, gspE, gspD, gspC, entD, fepA, fes, entF, fepE, fepC, fepG, fepD, ybdA, fepB, entC, entE, entB, entA, rcsB, fur, ompA |
| *Shigella flexneri* | phoP, csgC, csgA, csgB, fimB, fimE, fimA, fimI, fimC, fimF, fimG, fimH, ompA, fur, entA, entB, entE, fepB, entS, fepD, fepG, fepC, entF, fes, fepA, entD, acrB, rcsB, rhs, rpoS |
| *Vibrio cholerae* | ace, zot, ctxA, ctxB, rtxA, rtxC, vgrG-1, motY, motB, cqsA, flgM, cheV, cheR, flgB, flgC, flgD, flgE, flgF, flgG, flgH, flgI, flgJ, flgL, flaA, flaC, mshD, mshC, mshA, mshB, mshF, mshG, mshE, mshN, mshM, mshL, mshK, mshJ, mshI, mshH, hlyA, gbpA, vasH, cheW, cheB, cheA, cheZ, cheY, fliA, flhF, flhA, flhB, fliR, fliQ, fliP, fliO, fliN, fliM, fliL, fliK, fliJ, fliI, fliH, fliG, fliF, fliE, fliS, flaI, fliD, flaG, flaB, flaD, flaE, motX, epsN, epsM, epsL, epsK, epsJ, epsI, epsH, epsG, epsF, epsE, epsC, pilC, pilB, nanH, luxS, tcpI, tcpP, tcpH, tcpA, tcpB, tcpQ, tcpC, tcpR, tcpD, tcpS, tcpT, tcpE, tcpF, tcpJ, acfB, acfC, acfA |
| *Vibrio parahaemolyticus* | tlh, VPA0450, vpadF, vxsC, virG, exsA, exsD, vscB, vscC, vscD, vscF, vscG, vscH, vscI, vscJ, vscK, vscL, vopS, vopR, vecA, vopQ, vscU, vscT, vscS, vscR, vscQ, vscP, vscO, vscN, vopN, tyeA, vcr1, sycN, vcr2, vscX, vscY, vcrD, vcrR, vcrG, vcrV, vcrH, vopB, vopD, fliG, cheY, cheW, flgB, mam7 |
| *Clostridium difficile* | CD0873, cwp84, cwp66, fbpA, fbp68, iap, ibp, groEL, toxB, CD2831, zmp1 |
